# Luminescent ingestible electronic capsules for *in vivo* regulation of optogenetic engineered bacteria

**DOI:** 10.1101/2024.05.24.595681

**Authors:** Lianyue Li, Zhijie Feng, Xinyu Zhang, Mingshan Li, Haoyan Yang, Dawei Sun, Hongxiang Li, Huimin Xue, Hanxin Wang, Yaxin Wang, Le Liu, Yanyan Shi, Duo Liu, Taofeng Du, Hanjie Wang

## Abstract

The ideal engineered microbial smart-drug should be capable of functioning on demand at specific sites *in vivo*. However, precise regulation of engineered microorganisms poses challenges in the convoluted and elongated intestines. Despite the promising application potential of optogenetic regulation strategies based on light signals, the poor tissue penetration of light signals limits their application in large experimental animals. Given the rapid development of ingestible electronic capsules in recent years, taking advantage of them as regulatory devices to deliver light signals *in situ* to engineered bacteria within the intestines has become feasible. In this study, we established an electronic-microorganism signaling system, realized by two Bluetooth-controlled luminescent electronic capsules were designed. The “Manager” capsule is equipped with a photosensor to monitor the distribution of engineered bacteria and to activate the optogenetic function of the bacteria by emitting green light. The other capsule, “Locator”, can control the in situ photopolymerization of hydrogels in the intestines via ultraviolet light, aiding in the retention of engineered bacteria at specific sites. These two electronic capsules are expected to work synergistically to regulate the distribution and function of engineered bacteria *in vivo*, and their application in the treatment of colitis in pigs is currently being investigated, with relevant results to be updated subsequently.

## Introduction

Orally administered engineered probiotics perform functions such as secretion, metabolism, and sensing in the intestinal tract, making them a potential next-generation smart drug^1-3^. An ideal smart drug should be capable of functioning at designated locations in the gut as needed^4,5^. However, the convoluted and elongated nature of the intestines makes it challenging to spatiotemporally control the distribution and function of engineered probiotics after oral administration. Thus, crafting strategies to better control engineered microbes is crucial for their application advancement.

Currently, a variety of methods for regulation of the engineered microbes *in vivo* have been developed, including chemical molecules or physical signals^6-9^. Among these, light signals have the advantages of strong orthogonality and high spatiotemporal precision in vivo^10^. By constructing optogenetic engineered microbes, the functions of their equipped optogenetic circuits can be regulated by visible light. Optogenetic regulation strategies have already been applied in mouse models, including light-controlled engineered bacteria for treating colitis^11,12^, tumors^13,14^, et al. Besides, light signals can also control the cross-linking of photosensitive hydrogel materials in vivo^15,16^, allowing for the control of drug accumulation at lesion sites through the gel. Although light regulation strategies have clear advantages in application, their extremely poor tissue penetration making it impossible to deliver to deep tissues in a non-invasive manner^17^. To explore a non-invasive strategy to deliver light to the intestines is key to breaking through this limitation.

Recent advancements in ingestible electronic capsules offer a non-invasive, convenient way for intestinal operations without discomfort, akin to taking a regular pill^18,19^. The advancement in microelectronics has allowed for the creation of highly customizable electronic capsules. Ingestible electronic capsules can serve as alternatives to traditional endoscopies^20,21^ and treat diseases by stimulating the intestines or delivering medication, including for constipation^22^ or inappetency^23^. Based on this, using electronic capsules as a potential device presents a promising solution for delivering light within the intestines, and controlling the capsules from outside the body to apply light regulation signals to engineered probiotics.

After electronic capsules enter the intestines, accurately applying light control signals to the optogenetic engineered bacteria becomes extremely difficult due to challenges in determining the microbial distribution. Thus, an ideal electronic capsule should be capable of real-time monitoring of engineered microbial signals within the intestines and be able to controllably emit light to activate the bacteria^24,25^. This electronic capsule requires the inclusion of a bio-signal sensing module, Bluetooth signal transmission module, and a light excitation module in the electronic capsule, which places higher demands on the capsule’s size and battery storage. Moreover, given the non-directional distribution of orally administered engineered bacteria in the intestines, it is necessary for the electronic capsule to control the residency of engineered microbes in specific intestinal segments after oral administration. The electronic capsule devices should taking roles of both monitoring the engineered microbial signals and regulating these microbes designated to cluster at the lesion site to maximize therapeutic effects.

In this research, two types of bioelectronic capsules customized for intestinal tract applications were developed. The first “Manager” capsule, equipped with a photosensor, identifies the bacterial luciferase bioluminescence signals from the engineered bacteria to determine their location and emits green light to activate the microbial optogenetic functions. The second “Locator” capsule emits 365 nm light to achieve the photocrosslink of polyethylene glycol diacrylate (PEGDA) with engineered bacteria, thereby controlling microbial spatial distribution within the intestine. The concept of the using this electronic-microorganism system for regulating the functions of optogenetic engineered probiotics has been proved effective in the pig model.

**Schematic 1.**
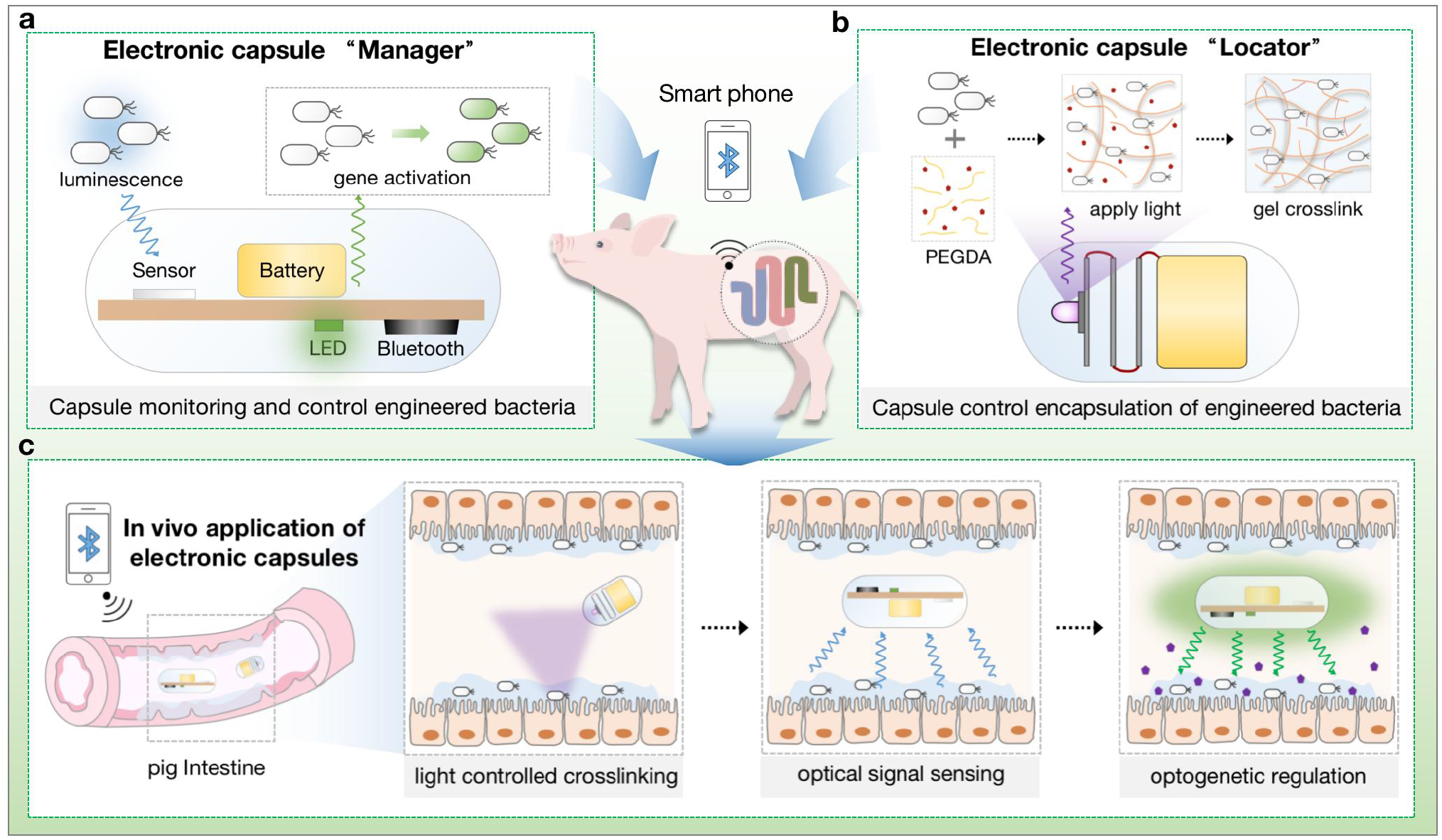
Ingestible luminescent electronic capsule for control of optogenetic engineered bacteria. **a**. The electronic capsule “Manager” monitors the chemical luminescence emitted by engineered bacteria and enables light-controlled regulation of their functionality. **b**. The electronic capsule “Locator” emits controllable ultraviolet light to control the crosslinking of hydrogel materials. **c**. The electronic capsule serves as a platform for the regulation of engineered bacteria in the pig intestine.

### Design and functional testing of electronic capsule “Manager”

To meet the requirements for monitoring and regulating the engineered bacteria, the capsule’s printed circuit board (PCB) was designed to include a 3×3 mm^2^ high-sensitivity photodetector, three 560 nm wavelength LEDs, and voltage conversion modules. Additionally, it comprises Bluetooth and an antenna for wireless connection to smartphones (**Fig. 1a** and Extended Data Fig. 1a). Powered by a button battery, the PCB is encased in a transparent plastic (PET) shell, which provides resistance to acid corrosion during the passage of the electronic capsule in stomach while allowing light transmission. The overall dimensions are 13 mm in diameter and 33 mm in length **(Fig. 1b)**. The developed control software enables searching for the capsule’s Bluetooth signal, connecting and controlling the functions of photosensor and the photoemitter LED within the interface, and displaying real-time electrical signal variations from the photosensor **(Fig. 1c)**. To perfectly match the electronic capsule functions, the biosafe *Escherichia coli* strain Nissle1917 (EcN)^26^ was engineered to constitutively express the luciferase *LuxCDABE* gene cluster, allowing it to continuously emit detectable light (EcN-Lux) (Extended Data Fig. 2a).

**Fig. 1.**
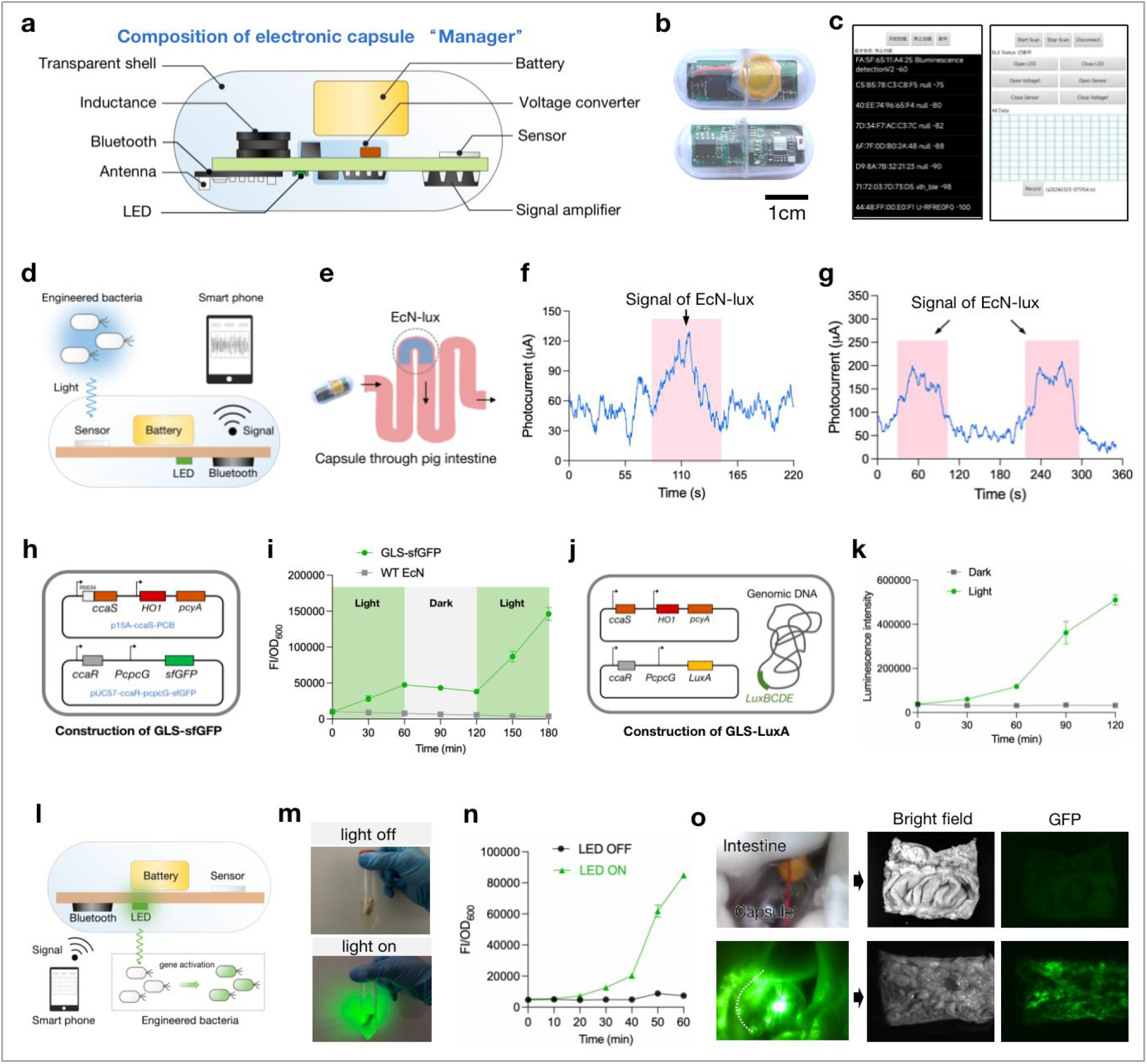
Design, functional testing and optimization of electronic capsule “Manager”. **a**. Composition of the “Manager” electronic capsule’s functional modules. **b**. Physical photograph of the electronic capsule, showing both sides of the electronic circuit board, bar=1cm. **c**. Connection and operation interface of the electronic capsule’s control software. After device search, the electronic capsule’s device name can be found on the left interface; clicking on it will switch to the right interface, where functions can be controlled on and off with buttons. **d**. The electronic capsule monitors the chemiluminescent signal of engineered bacteria and transmits photocurrent values to the smartphone. **e**. The electronic capsule slowly passes through a segment of pig intestine, where 2mL of engineered bacteria EcN-Lux with OD600=1 was pre-added at a specific location, and the capsule records photocurrent values throughout the intestine. **f**. Signal values detected by the electronic capsule in the intestinal lumen with a distribution of engineered bacteria at one site, where the peak in the pink area represents the detected signal of bacteria distribution. The data has been smoothed, with raw data shown in Extend. **g**. Photocurrent signals recorded by the electronic capsule when there are two distributions of engineered bacteria in the intestinal lumen. **h**. The engineered bacterial gene circuit for sfGFP expression in response to green light consists of two functional plasmids: p15A-ccaS-PCB and pUC57-ccaR-pcpcG-sfGFP. **i**. GLS-sfGFP can initiate and cease expression of the reporter gene sfGFP in response to switching between light and dark states. During induction of the engineered bacteria, a 2-hour green light exposure is applied, followed by 2 hours of darkness, and then another 2-hour light exposure. The induction efficacy is assessed by the ratio of green fluorescence intensity to OD600. **j**. The introduction of green-light optogenetic circuit to the GLS-LuxA strain makes it emit chemiluminescence in response to green light, involves integration of the *luxCDBE* genes from the *lux* operon into the genome of the engineered bacteria. Expression of the *luxA* gene is controlled by green light, enabling optogenetic regulation of chemiluminescence in the engineered bacteria. **k**. The optogenetically regulated engineered bacteria GLS-LuxA exhibit chemiluminescent functionality in response to green light control. **l**. The electronic capsule can emit controllable green light to activate the function of green light sensitive (GLS) optogenetically engineered bacteria GLS-sfGFP. **m**. Off and on states of the electronic capsule’s LED in the bacterial liquid. **n**. Samples are taken every 10 minutes from engineered bacteria in both dark and illuminated states to test green fluorescence intensity (FI) and OD600 values. The activation status of engineered bacteria GLS-sfGFP is evaluated through the ratio of FI to OD600. **o**. Validation of the electronic capsule’s effect in activating engineered bacteria within the pig colon. The state of the capsule in the pig colon is captured via endoscopy, and after illumination, the opened pig colon is imaged using a small animal in vivo imaging system.

**Fig. 2.**
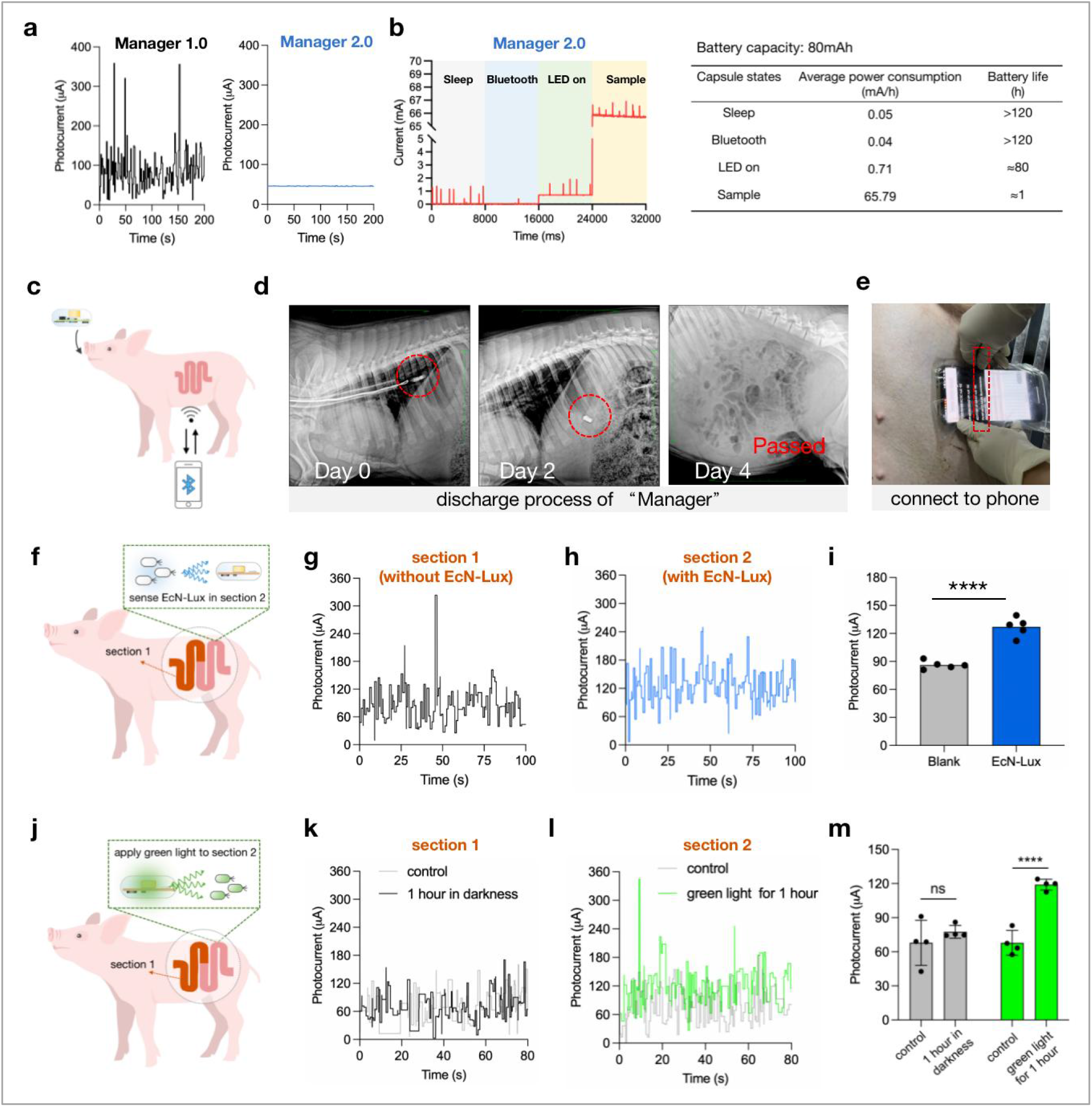
In vivo testing of the monitoring and regulation functions of “Manager” capsule. **a**. Comparison of photocurrent values of blank samples tested with version 1.0 and 2.0 of the electronic capsule, where neither group of signals has been post-processed. b. Tests of current consumption values of the electronic capsule in different states, and battery life tests with different functions activated. “Sleep” refers to the state where Bluetooth is not connected but the capsule device can be searched, “Bluetooth” refers to the state where the capsule’s Bluetooth is connected, “LED on” indicates all three LEDs of the electronic capsule are activated, and “Sample” refers to the capsule’s sensor working and continuously transmitting photocurrent values. **c**. The electronic capsule “Manager” is orally administered and controlled via Bluetooth connection through a mobile phone. **d**. X-ray imaging of the capsule inside the pig on days 0, 2, and 4, with the capsule’s position indicated by red circles. **e**. The Bluetooth signal of the capsule inside the pig’s body can be detected through the control software. Clicking on the capsule device name will lead to the connected control interface. **f**. Two equally sized segments were delineated within the pig’s intestine to investigate the in vivo monitoring capability of the capsule “Manager” for engineered bacteria signals. **g**. The signal waveforms from the segment 1, and the segment 1 underwent no manipulation. **h**. The signal waveforms from the segment 1, and the segment 2 received EcN-Lux. The position of the capsule was controlled by the length inserted via endoscopy. **i**. Comparison of the mean signal values in different segments. **j**. Similarly, investigating the optogenetic control effects of the electronic capsule on engineered bacteria in different segments of the intestine. **k**. The signal waveforms before and after operations in the segment 1. The segment 1 received GLS-LuxA and kept dark. **l**. The signal waveforms before and after operations in the segment 1. The segment 2 received GLS-LuxA and followed by 2 hours of light exposure. **m**. Calculate the mean values of the signals from the two segments in Fig. 2k and Fig. 2l.

To test the particular microbial sensing function of the capsule, it was employed to detect the engineered microbial cells with different luminescence intensities **(Fig. 1d)**. Changes in the capsule’s photocurrent were recorded, showing a positive trend correlating with results from an luminometer (Extended Data Fig. 2b). The detection capability boundary of the “Manager” capsule was determined by testing the engineered strains with intensity from 10^5^ to 10^6^ CFU (or OD600 from 0.1 to 1), showing the capability of detecting as low as about 10^5^ CFU of strains and the bioluminesence signals emitted from the strains as far as 1.5 cm away (Extended Data Fig. 2c). *In vitro*, a pig colon was used to simulate the capsule’s working environment; when wild-type EcN or EcN-Lux was added to the intestinal lumen, the photocurrent values from the electronic capsule could effectively differentiate samples containing EcN-Lux (Extended Data Fig. 3a-c). Further, the engineered strains were artificially enriched at one or two positions. As the capsule slowly passed through the lumen, the changes in photocurrent reflected peaks corresponding to the locations of the engineered bacteria, demonstrating the electronic capsule’s capability to monitor the gathering of the engineered strains in intestines **(Fig. 1e-g)**.

**Fig. 3.**
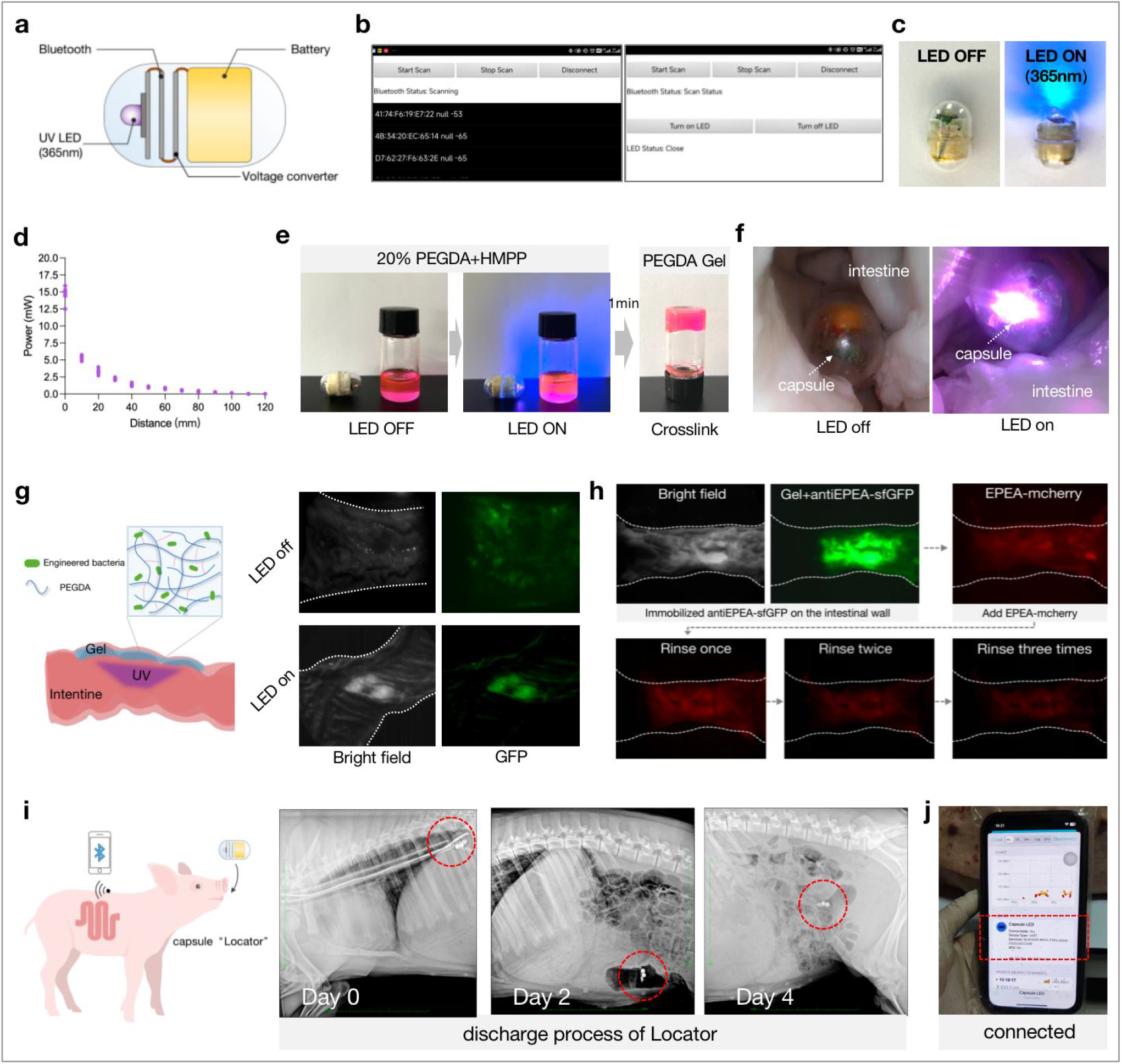
Design and in vitro functional validation of the “Locator” electronic capsule. **a**. The composition of the electronic capsule “Locator” includes ultraviolet LED, Bluetooth module, battery, and voltage conversion module. **b**. The control software interface of the electronic capsule supports Bluetooth search connection and functional operation. **c**. The electronic capsule is capable of controlled luminescence, with the same shell material and encapsulation method as “Manager”. **d**. The intensity of ultraviolet light emitted by the electronic capsule at different distances. **e**. The electronic capsule controls the crosslinking of PEGDA material. The hydrogel itself is transparent, and red dye Rhodamine is added for easy observation. **f**. In vitro simulation of the electronic capsule’s controllable crosslinking hydrogel in the pig intestine. The image shows engineered bacteria luminescing in the intestine, captured by a handheld endoscope. **g**. PEGDA is mixed with engineered bacteria expressing sfGFP (EcN-sfGFP) and introduced into the pig intestine. Luminescence of the electronic capsule is controlled to immobilize the bacteria on the intestinal wall, with green fluorescence indicating the distribution of engineered bacteria. **h**. EPEA-mcherry can adhere to the gel regions encapsulating antiEPEA-sfGFP. Green represents gel regions, while red represents EPEA-mcherry. After three washes, the red fluorescence at the gel site remained, indicating the binding of EPEA-mCherry to the hydrogel. i. X-ray images of the position of the electronic capsule “Locator” in the pig intestines on days 0, 2, and 4 after oral administration. j. The capsule signal in the pig intestines can be detected externally via mobile phone Bluetooth connectivity.

To equip the engineered strains with optogenetic regulatory function, two-component ccaS/ccaR^27^ green light regulatory circuit was constructed in the strain for controlling the expression of the reporter gene sfGFP, getting the strain GLS-sfGFP **(Fig. 1h)**. Compared to the blue light regulatory system previously employed by our group, the green light system demonstrated a higher induction fold (Extended Data Fig. 4a-c). Through optimization of plasmid copy number and RBS, the relative fluorescence intensity of the reporter gene in the engineered bacteria was upregulated approximately 12.5-fold after two hours of green light exposure (Extended Data Fig. 4d-f). The engineered bacteria GLS-sfGFP were able to switch between light and dark states in response to illumination **(Fig. 1i)**. To facilitate in vivo characterization of the engineered bacteria activation, a strain GLS-LuxA, capable of chemiluminescence after green light stimulation, was engineered and tested **(Fig. 1j-k)**. Preliminary validation of green light activation for in vivo activation of optogenetically engineered bacteria was conducted using GLS-LuxA in the mouse intestine (Extended Data Fig. 4g-h).

**Fig. 4.**
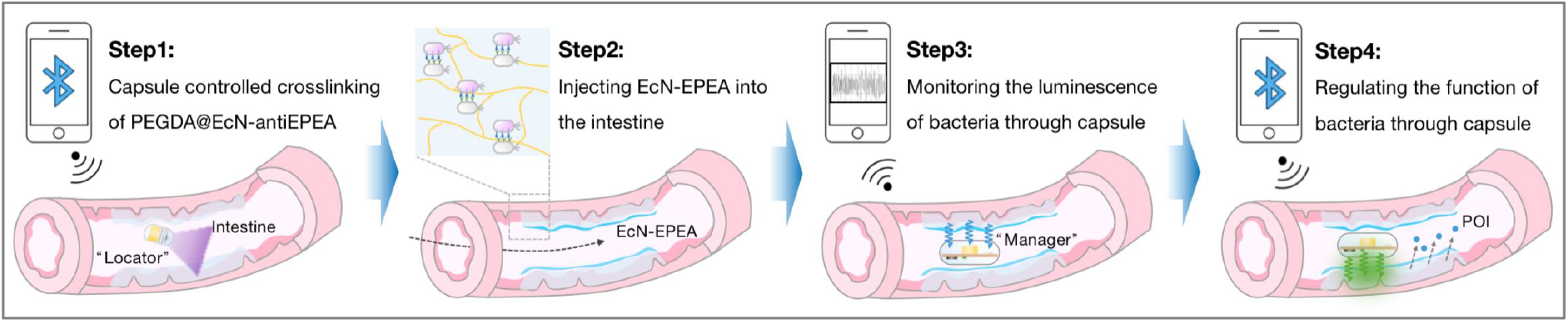
Using dual capsules in tandem as an engineered bacterial gut regulation platform. Operational steps of the dual-capsule engineered bacterial regulation platform.

Subsequent tests were conducted on the electronic capsule’s ability to regulate optogenetic engineered strains through the photoemitting LED **(Fig. 1l)**. The capsule’s light emission within the bacterial liquid was controlled to test the relative green fluorescence intensity of the liquid illuminated for different durations **(Fig. 1m)**. After one hour of illumination by the capsule, the fluorescence intensity of the microbial liquid increased by about 10-fold, indicating the capsule’s capacity for rapid functional regulation of the engineered strain **(Fig. 1n)**. When green light-responsive engineered bacteria were added to the lumen of the pig colon and the electronic capsule was controlled to illuminate within the lumen, widespread green fluorescence could be detected on the intestinal wall after one hour of illumination, demonstrating the capsule’s capability of activating the engineered bacteria within 1 h **(Fig. 1o)**.

### In vivo testing of the monitoring and regulation functions of “Manager” capsule

To ensure that the capsule met the requirements for *in vivo* applications, optimizations were made to the capsule’s signal and energy consumption, leading to the design of the electronic capsule “Manager” version 2.0 (Extended Data Fig. 1b). Compared to the first version, the photocurrent noise in version 2.0 was effectively reduced **(Fig. 2a)**. Additionally, the continuous energy consumption of high-energy modules was avoided for the purpose of enabling the separate controllability of the capsule’s Bluetooth, LED, and sensor modules,. Controlling the activation of specific functions as needed allowed the capsule’s battery working life to far exceed to cover the time period from oral administration to excretion of the “Manager” capsule **(Fig. 2b)**.

To prove that the electronic capsule “Manager” capsule could monitor and regulate the cellular functions of the engineered strains in live intestine, pigs were chosen as experimental animals because their abdominal tissue thickness and intestinal size are far more similar to humans **(Fig. 2c)**. In an initial experiments, the electronic capsule was orally administered to the anesthetized pigs and the pig swallowed it voluntarily. X-ray imaging was used to monitor the excretion process of the electronic capsule (Extended Data Fig. 5a). After excretion, the capsule shell was found to be damaged due to chewing in the pig’s mouth (Extended Data Fig. 5b). Therefore, the administration method of the capsule was optimized by inserting the electronic capsule into the front end of a hollow, flexible rubber tube approximately 50 cm long with an inner diameter of 1cm (Extended Data Fig. 5c). Similar to a gastroscope, the flexible tube was inserted into the stomach, and the capsule was pushed out of the rubber tube using a metal wire, thus avoiding the damage of chewing force to the capsule (Extended Data Fig. 5d).

After introducing the electronic capsule into the pig’s stomach, the excretion process of the electronic capsule was monitored **(Fig. 2d)**. X-ray imaging of the pig’s abdomen was conducted on days 0, 2, and 4, allowing visualization of signals generated by the metallic components within the electronic capsule. The time it takes for the electronic capsule to be expelled depended on various factors such as the time taken to enter the intestine from the stomach and the pig’s food intake. Generally, it enters the intestine on the second day after ingestion and is expelled on the fourth day. During the time period of the capsule movement in the intestine, its Bluetooth signal could be detected externally through an mobile phone application (APP) **(Fig. 2e)**, allowing connection and controlling of the electronic capsule’s functions, such as turning them on or off (Extended Movie S1).

As mentioned above, the initial approach involved orally administion of both the capsule and the engineered strain, but it did not yield the desired results. One of the probable reasons was the destruction of the engineered bacteria by gastric acid. Additionally, the longer excretion period and the considerable length of the intestine made it difficult to accurately determine the capsule’s position. Considering these factors, a compromise experimental approach was adopted. The electronic capsule was affixed to the front end of an endoscope and introduced through the pig’s anus into the intestine, simulating its actual usage scenario (Extended Data Fig. 6). This compromised approach aimed to reduce uncontrollable factors during the experiment and did not harm the pig’s intestine.

In the pig’s intestine, two segments were delineated in the same pig model, each with a length of about 20 cm. The localization of the capsule could be controlled in each respective segment, and engineered strains could be delivered to specific segments through the anus **(Fig. 2f)**. Then, EcN-Lux was introduced into segment 2, and signals of two segments were monitored externally **(Fig. 2g-h)**. The average signal of segment 2 was significantly higher than the segment 1, demonstrating the electronic capsule’s ability to monitor engineered bacteria signals in vivo **(Fig. 2i)**. Subsequently, equal amounts of GLS-LuxA strains were introduced into both segment 1 and segment 2. The segment 2 was exposed to 2 hours of light through the capsule **(Fig. 2j)**. Signal values from all segments were then tested **(Fig. 2k-l)**, and the average signal from segment 2 after light stimulation was significantly higher than the dark segment 1, proving the electronic capsule’s ability to regulate engineered bacteria in vivo through light exposure **(Fig. 2m)**.

### Design and functional validation of the “Locator” electronic capsule

Ensuring the presence of the engineered bacteria at specific locations within the intestinal tract presents a challenge due to their random distribution upon application. Concentrating these bacteria at the intended sites can significantly enhance therapeutic efficacy. Photo crosslinking of the hydrogels, such as PEGDA-based hydrogels, have shown promise in immobilizing cells at wound sites^28^. Thus, utilizing light-controllable hydrogels to crosslink at targeted locations within the intestine offers a promising approach to regulate the distribution of the engineered bacteria.

To achieve controllable crosslinking of hydrogels within the intestine, we designed an electronic capsule named “Locator” **(Fig 3a)**. This capsule, connectable via Bluetooth, regulated the UV LED switch to emit ultraviolet light for in-situ crosslinking of hydrogels within the intestinal tract **(Fig 3b-c)**. Enclosed in packaging akin to the “Manager”, the capsule measured approximately 10 mm in diameter and 15 mm in length (Extended Data Fig. 7a). The UV light power decays with distance, peaking at around 15 mW **(Fig 3d)**. Under the capsule’s UV irradiation, PEGDA hydrogels could crosslink in about 1 min **(Fig 3e)**. The crosslinking time of the hydrogel was correlated with the distance from the capsule to the gel, as the capsule can complete crosslinking of gel located 5 cm away in approximately 15 minutes, demonstrating its ability to crosslink a wide range of gels (Extended Data Fig. 7b). Additionally, temperature variations during the 15 minutes operation of the electronic capsule *in vitro* were proved, not to adversely affect the intestines (Extended Data Fig. 7c). In vitro simulation of hydrogel crosslinking within the intestinal tract was conducted using an *in vitro* porcine intestine. The capsule could be controlled via a mobile phone to emit light in the pig intestines **(Fig 3f)**. PEGDA solution mixed with photosensitizer was applied to the inner wall of the pig intestines. By controlling the capsule to illuminate the intestines for 15 minutes, the gel was crosslinked onto the intestinal wall. After cutting open the intestines and lifting them vertically, the PEGDA solution in the intestines without illumination flowed away (Extended Data Fig. 8a). The precursors of the hydrogel were mixed with engineered bacteria EcN-sfGFP and applied to the inner wall of the intestine. By illuminating with the capsule, the engineered bacteria could be immobilized at specific locations along the intestinal segment **(Fig 3g)**. The gel material and crosslinking process did not affect the activity of the strains (Extended Data Fig. 8b).

When taking efforts to integrate the roles of “Locator” capsule with the “Manager” capsule, we primarily tried to immobilize the strain EcN-Lux by using the photocrosslinking hydrogel. But the strains embedded in the gel failed to emit the expected chemiluminesce of bacterial luciferase, probably due to interference from the photocosslinking hydrogel monomers with luciferase catalyzing function (Extended Data Fig. 9a). Therefore, we decided to strip out the strains emitting luminescence from the photocorsslinking gels and made the strains and the gels converge by particular affinity binding forces. Previous studies have shown that specific bacterial binding can be achieved through the display of complementary proteins antiEPEA and EPEA on their surfaces^29^, respectively (Extended Data Fig. 9b). Leveraging this mechanism, we encapsulated bacteria displaying antiEPEA within the hydrogel and externally introduced bacteria expressing mCherry along with surface-displayed EPEA. Through protein-protein interaction, EPEA-mCherry exhibited stronger binding affinity to the hydrogel (Extended Data Fig. 9c). Subsequent in vitro validation using pig intestines demonstrated the improved attachment of EPEA-mCherry to the gel-coated areas of the intestinal wall **(Fig 3h)**. This way allowed the attachment of chemiluminescent bacterial strains to the hydrogel surface without significantly affecting the chemiluminescence (Extended Data Fig. 9d). The hydrogel encapsulating antiEPEA strains could serve as targets within the intestine. Subsequently introduced EPEA-Lux bacteria were expected to distribute more abundantly in the gel-coated areas, potentially enabling the “Manager” to monitor signals more effectively. The excretion process of the “Locator” capsule post-oral administration was monitored using X-ray imaging **(Fig 3i)**. Simultaneously, the connection to and functional control of the capsule’s Bluetooth signal were achieved from outside the body **(Fig 3j)**.

### Employment of dual capsules in tandem as a controllable probiotic functionality platform

After validating the functions of the two electronic capsules separately, the possibility of their combined use was further considered. A schematic of this approach was shown in the figure **(Fig 4)**. In an ideal application scenario, the process might begin with the oral administration of a mixture of EcN-antiEPEA and the photopolymerizable agent PEGDA, with crosslinking controlled by the electronic capsule “Locator”. Subsequently, the engineered strain GLS-Lux, displaying EPEA on their surface, would be orally administered to allow specific binding between these two groups of strains via surface protein affinity. Finally, the electronic capsule “Manager” would be administered to monitor microbial signals and regulate the optogenetic function of the engineered bacteria by emitting green light.

To demonstrate the in vivo application effectiveness of the dual-capsule regulation platform, relevant experiments in live pig intestine were underway. These included exploring the use of the dual-capsule light-regulation strategy for treating colitis in pigs. Colitis was induced in pigs by oral administration of DSS, and meanwhile a healthy group of pigs were not treated by DSS as a positive control group. The control group of pigs received daily oral administration of a strain capable of light-induced expression of porcine IL-10 protein (GLS-IL), and the treatment group used the dual-capsule regulation platform to activate the therapeutic function of the GLS-IL engineered bacteria via light control. The entire treatment period spanned two weeks, during which macro indicators such as stool condition and intestinal temperature were monitored. Fecal and blood molecular markers related to colitis were tested, and intestinal tissue sections were analyzed for pathological features. The therapeutic effect on pig colitis was evaluated through multidimensional data. These experiments are ongoing, and results will be published in subsequent versions.

## Discussion

This research proposes a novel in vivo optogenetic regulation system for engineered gut bacteria using ingestible electronic capsules controlled via Bluetooth. The system ensures precise light delivery to bacteria within the intestines. Two distinct capsules were designed to address positioning and functionality issues. The “Manager” capsule monitors luciferase-expressing bacteria using a photosensor and activates bacterial functions with green light. The “Locator” capsule emits ultraviolet light to crosslink photosensitive hydrogels, aiding bacteria retention at specific intestinal sites. The engineered bacterial in vivo regulation system with ingestible electronic capsules offers key advantages. First, it provides visibility: unlike traditional methods that struggle with uncertain bacterial distribution in the intestines, the “Manager” capsule can monitor bacterial luminescence to accurately apply light signals. Second, it overcomes tissue penetration issues by delivering light directly in situ within the intestines, making the process more efficient. Third, it is easy to operate: the capsules connect to a smartphone via Bluetooth, with user-friendly software to receive and send signals, allowing for simple home management.

Engineered bacteria represent a promising new generation of smart therapeutics, and precise in vivo regulation is crucial for achieving precision medicine and clinical applications. Previous in vivo regulation efforts have been confined to small animal models, with applications in larger animals (e.g., pigs, monkeys) hampered by the limited tissue penetration of regulatory signals. This study integrates electronic and material sciences to develop a feasible strategy for regulating engineered bacteria in the pig intestine. Ingestible electronic capsules emit light signals in situ, directly targeting intestinal bacterial strains or photopolymerizable materials, allowing for immediate execution of external commands. The proposed framework, “engineered bacteria + electronic devices + functional materials”, offers a generalizable approach. Electronic devices serve as interfaces for external communication, while functionally designed bacteria and materials act as functional modules upon receiving signals from the electronic devices. This concept could broaden the scope of in vivo applications. For example, by using electronic devices to monitor additional physiological indicators within the intestines, the system can assist in determining disease states and whether activation of the therapeutic function is needed. By incorporating photosensitive therapeutic nanomaterials, the system can offer more options for the disease treatment module. The notion of “semiconductor synthetic biology” has already been introduced, and this study extends that concept, providing new possibilities for the interaction between devices and bacterial strains.

The proposed approach has been preliminarily validated both in vitro and within the pig intestine. But several challenges have emerged as the work progresses, including: (1) Limited in vivo characterization techniques for large animals, with areas of the intestine that are inaccessible to endoscopy, making it difficult to directly observe or test the intestinal response of materials and bacterial strains. (2) The complexity of the digestive system in large animals is significantly higher than that in mice, and the impact of digestive fluids on orally administered strains and materials far exceeds the values observed in mouse experiments, necessitating considerations for protecting strains or materials post-ingestion. (3) The size and battery life of the electronic capsules currently only meet the basic experimental requirements; further optimization of battery life and size is needed to meet clinical trial standards. Presently, the research has progressed to exploring the effects of the combined use of the two capsules, including validating this strategy in a pig colitis model. Future research outcomes will provide further insights into the practical application of this approach. Please stay tuned for upcoming results.

In summary, the construction of this new “electronic capsule-optogenetic engineered bacteria” gut regulation system holds significant potential for advancing the clinical application of engineered bacteria. It promises to facilitate personalized medicine based on living engineered bacterial therapeutics.

## Notes

### Competing Interest Statement

The authors have declared no competing interest.

